# Nano-organization of synapses defines synaptic release properties at cortical neuron dendritic spines

**DOI:** 10.1101/2025.02.13.637710

**Authors:** Haani Jafri, Samantha J. Thomas, Sung Hoon Yang, Rachel E. Cain, Matthew B Dalva

## Abstract

Visualization of the submicron organization of excitatory synapses has revealed an unexpectedly ordered architecture consisting of nanocolumns of synaptic proteins that group into nanomodules which scale in number as spine size increases. How these features are related to synaptic function has remained unclear. Here, using super-resolution followed by live-cell line-scan imaging, we find that the size of the smallest miniature calcium and glutamate events are the same, regardless of whether spines have one or two nanopuncta of PSD-95, and that miniature synaptic response in all spines are best fit by a three term Poisson. Two nanomodule spines exhibit more large events without a significant change in event frequency, with the number of the largest events increasing disproportionately. These data support a model where nanomodules define sites of synaptic release and where the nanoarchitecture of synaptic proteins specifies subtypes of excitatory synapses, with increasing numbers of nanomodules increasing coordinated multivesicular release.

## Introduction

Synaptic connections underlie neuronal communication and memory in the nervous system and are implicated in many neurological diseases. Synaptic size varies and is linked to the strength of postsynaptic responses, with increasing spine size associated with increased synaptic strength^1,2^. The nanoscale organization of the synapse is also dynamic and linked to the size of the dendritic spine. The functional components of excitatory synapses – including ionotropic and metabotropic glutamate receptors, scaffolding proteins, and active zone proteins– organize into nanoclusters that align across the synaptic cleft forming trans-synaptic nanocolumns which optimize neurotransmission ^3–5^. These nanocolumns scale in number, but not size, as spine size increases, suggesting that they are modular structures (‘nanomodules’) that may relate to the function of the spine ^6^. Consistent with this model, following structural plasticity, which enhances synaptic strength and drives spine enlargement, the number of nanomodules increases in spines that undergo sustained increases in size^6,7^. This suggests that the nanoscale features of spines may specify differences in how synapses at larger and smaller spines function. However, it remains unclear whether the number of nanomodules at a spine is related to synaptic function or transmission.

To determine the relationship between synaptic nanomodules and properties of synaptic function, the nanoscale features within individual spines must be related to their synaptic responses. Electrophysiology techniques have a limited ability to record from single spines, because of their small size, and the difficulty of locating the source of synaptic inputs from recordings in the soma or dendrite ^8–11^. Additionally, voltage compartments, like the spines themselves, may alter signal propagation within the neuron ^12^. To overcome these limitations, we conducted live-cell STED nanoscopy followed by time-lapse imaging of single spines in neurons transfected with the calcium or glutamate sensors, GCaMP8f and iGluSnFR3, respectively. We further simplified the postsynaptic response by measuring amplitude and frequency of spontaneous miniature postsynaptic calcium or glutamate events under conditions where action potentials are blocked. Quantal analysis of these data indicates that one and two nanomodule spines both have the same size single quantal events but larger two nanomodule spines have more high amplitude multi-quantal events. These data suggest that the nanoscale organization of spines defines distinct excitatory synapse subtypes with distinct patterns of quantal spontaneous release.

## Results

### Observed miniature synaptic Ca^2+^ transients are NMDAR-dependent

Larger spines generate larger postsynaptic responses ^1,2^. However, how the nanoscale organization of the spine relates to single, spontaneous synaptic events has yet to be determined. To determine this relationship, we imaged the postsynaptic activity of spines with known nanostructure by co-transfecting neurons with mTurq2-tagged PSD95-FingR and GCaMP8f, which enabled the identification of dendritic spines with either one or two nanoclusters of PSD-95 using STED imaging (Fig. 1a, b, Supplementary Fig. 1a-c). The PSD95-FingR nanostructure of the spine was imaged using live-cell gated-STED and then the same area of dendrite was imaged with live-cell time-lapse confocal imaging (83 Hz, 64 x 64 pixels, 406 nm x 406nm pixel size) of single spines. STED imaging was reserved for nanomodule identification to reduce photodamage to neurons^13^. To visualize the relationship of PSD-95 nanostructure to neuronal activity, spontaneous NMDAR-dependent calcium transients were analyzed, which ensured that the post synaptic responses were not saturated. To isolate NMDAR activity, a cocktail of drugs was used to block non-specific sources of calcium, including activity-dependent potentials, intracellular calcium stores, AMPA-type glutamate receptors and metabotropic glutamate receptors (2 μm TTX, 40 μm nifedipine, 30 μm dantrolene, 10 μm LY341495, 20 μm NBQX, and 500 μm S-MCPG), allowing for NMDAR-dependent miniature synaptic calcium transients (mSCTs) to be quantified^14–16^. With these blockers in place, spontaneous activity does not saturate the NMDAR response, permitting quantification of NMDAR responses under magnesium-free conditions^17–19^. For imaging of glutamate, only action potentials were blocked.

**Figure 1:**
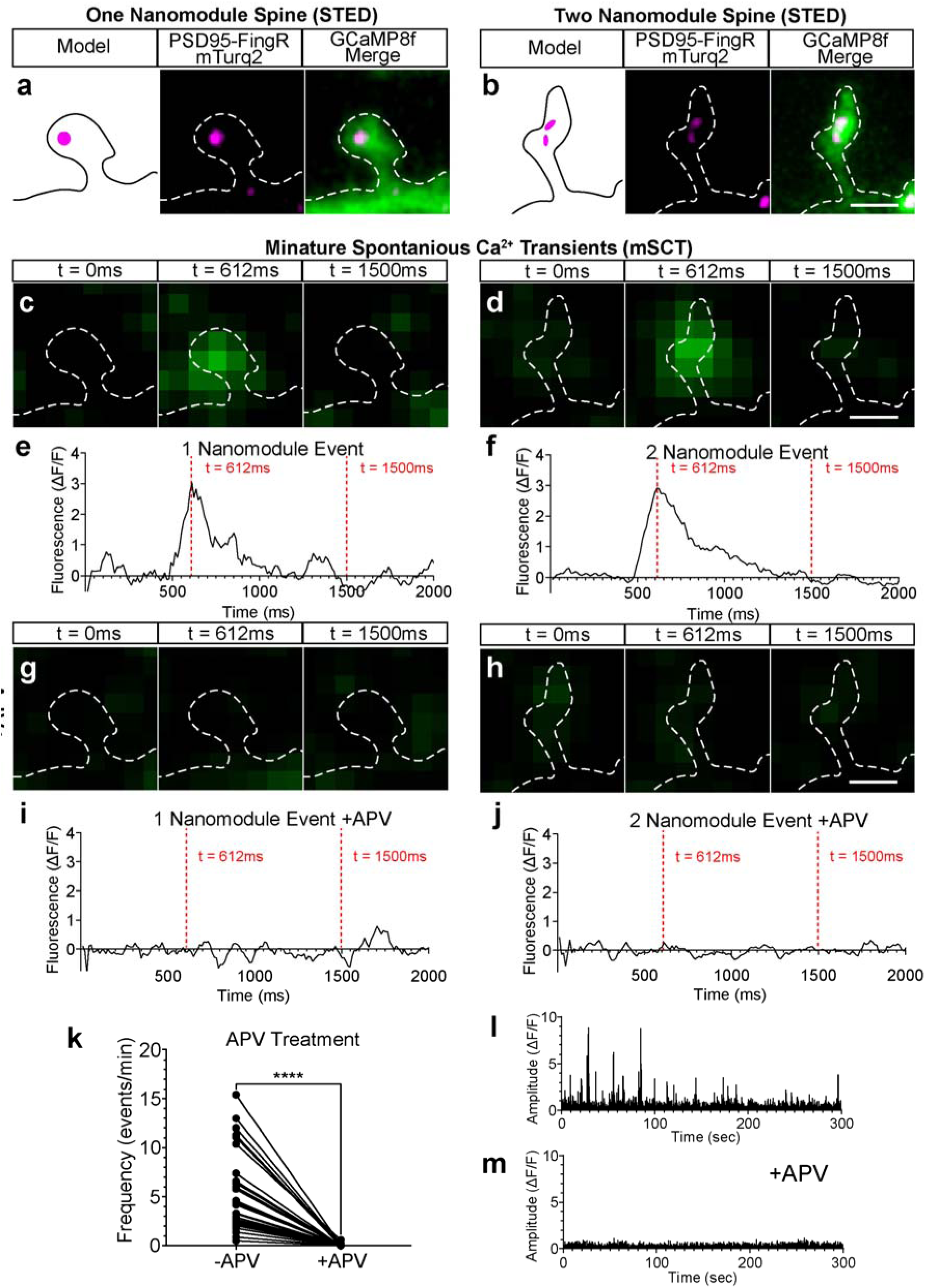
STED and Ca2+ imaging measurement of NMDAR-dependent events. **(a,b)** Schematic of live-cell STED nanoscopy of exemplary one (left) and two (right) nanomodule spines with live-cell STED resolved PSD-95 FingR (magenta) and GCaMP8f as a cell-fill (green). White dashed line shows morphology. Scale bar is 1μm. **(c,d)** The same one and two nanomodule spines at three different time points of a Ca^2+^-transient recorded under activity blockade: baseline before peak at t = 0ms (left panel), peak amplitude t = 612ms (middle panel), baseline after peak at t = 1500ms (right panel). Spine outlines (white dashed lines) are based on high pixel format imaging using STED. **(e,f)** Quantification of the same events with the time points labeled (red dashed lines). **(g-j)** Images and activity traces from the same spines treated with APV, demonstrating sustained baseline fluorescence. **(k)** Quantification of the frequency of spontaneous Ca^2+^-transients in individual dendritic spines before and after APV addition. N = 29 spines (**** p < 0.0001, paired t-test). **(l)** ΔF/F normalized fluorescence intensity trace of a dendritic spine under non-NMDAR activity blockade showing five minutes of spontaneous Ca^2+^- transient recording. **(m)** Same spine as shown in (l) following the addition of APV.

Calcium imaging revealed transient increases in fluorescence limited to the spine head (Fig. 1c, d, Movies S1, S2). Analysis of these calcium transients revealed a distinctive profile characterized by a rapid rising phase, a slower decay phase, and a subsecond time course, indicative of spontaneous NMDAR events^16,20–22^ (Fig. 1e, f). To confirm the NMDAR-mediated dependence of these events, neurons were treated with the selective NMDAR-antagonist (2R)-amino-5-phosphonovaleric acid (100 μm AP5). AP5 treatment blocked all fluorescence transients without changing the baseline fluorescence (Fig. 1g-m, Movies S3, S4). These data indicated that the recorded events were due to the activity-independent activation of NMDARs, likely via spontaneous vesicle fusion represented as mSCTs.

Miniature NMDAR-dependent synaptic transients exhibit a time course of approximately 200 milliseconds^23^. To determine the sensitivity of single spine imaging of GCaMP, the kinetics of mSCTs were recorded using different calcium sensors, GCaMP7f and GCaMP8f using 83 Hz 2D field scanning. (GCaMP6f did not resolve these events well, (data not shown), but see^16^.

GCaMP8f event kinetics were also recorded at either 83 Hz 2D field scanning and 4000 Hz 1D line scanning (Supplementary Fig. 2a-c). Consistent with design of GCaMP8f^24^, mSCTs recorded using GCaMP8f exhibited significantly faster rise times, showed significantly narrower full width at half maximum amplitude (FWHM), and had similar decay times as those recorded with GCaMP7f (Supplementary Fig. 2e, f). Moreover, line scans of GCaMP8f mSCTs demonstrated significantly faster rise times compared to 2D-scanned mSCTs (Supplementary Fig. 2d) without significant bleaching (Supplementary Fig. 3a), enabling visualization of the ∼200ms time course of NMDAR transients. Therefore, we used an approach combining STED imaging of PSD95-FingR with line scans of single spines filled with GCaMP8f to best capture spontaneous calcium events from spines with identified nanoscale organization.

### Visualization of mSCTs in individual spines reveals quantal event distributions

Larger spines generate larger postsynaptic responses and have more nanoclusters of pre- and postsynaptic proteins^6,7^. These clusters of pre- and postsynaptic proteins form in trans-synaptically paired nanomodules, or aligned nanocolumns, suggesting that synaptic nanomodules might be related to synaptic function^3,4,25^. The excitatory synapses found at dendritic spines transmit information by releasing glutamate from the presynaptic terminal, activating postsynaptic glutamate receptors. There are two ways that increasing the number of nanocolumns or nanomodules might alter the postsynaptic response. If the release of a single vesicle activates all postsynaptic nanopuncta, the size of the postsynaptic response should increase as more nanomodules are added. Alternatively, if a single vesicle can only activate the proximal nanopuncta, then the size of the postsynaptic response would not change with increasing number of nanomodules. These possibilities are explored below.

To explore the number of vesicles at spines with known nanoarchitecture, the nanoscale organization of spines was first determined by STED imaging, and then 4000 Hz line scan imaging of NMDAR-dependent mSCTs in single spines with known nanoscale organization was conducted (Fig. 2a). Spines expressing FingR were identified, and STED was used to determine whether they contained one or two PSD-95 nanomodules (Fig. 2b, d, l, n). These same spines were imaged for 10 or 20 minutes at 4000 Hz, where unitary mSCTs were both clearly visible (Fig. 2c, e, m, o, Movies S5-S8) and quantifiable (Fig. 2f, i, p, s) from the line scans. Only one and two nanopuncta spines were selected for 4000 Hz imaging because spines with larger numbers of nanopuncta are rare^6^ (<15% of all spines).

**Figure 2:**
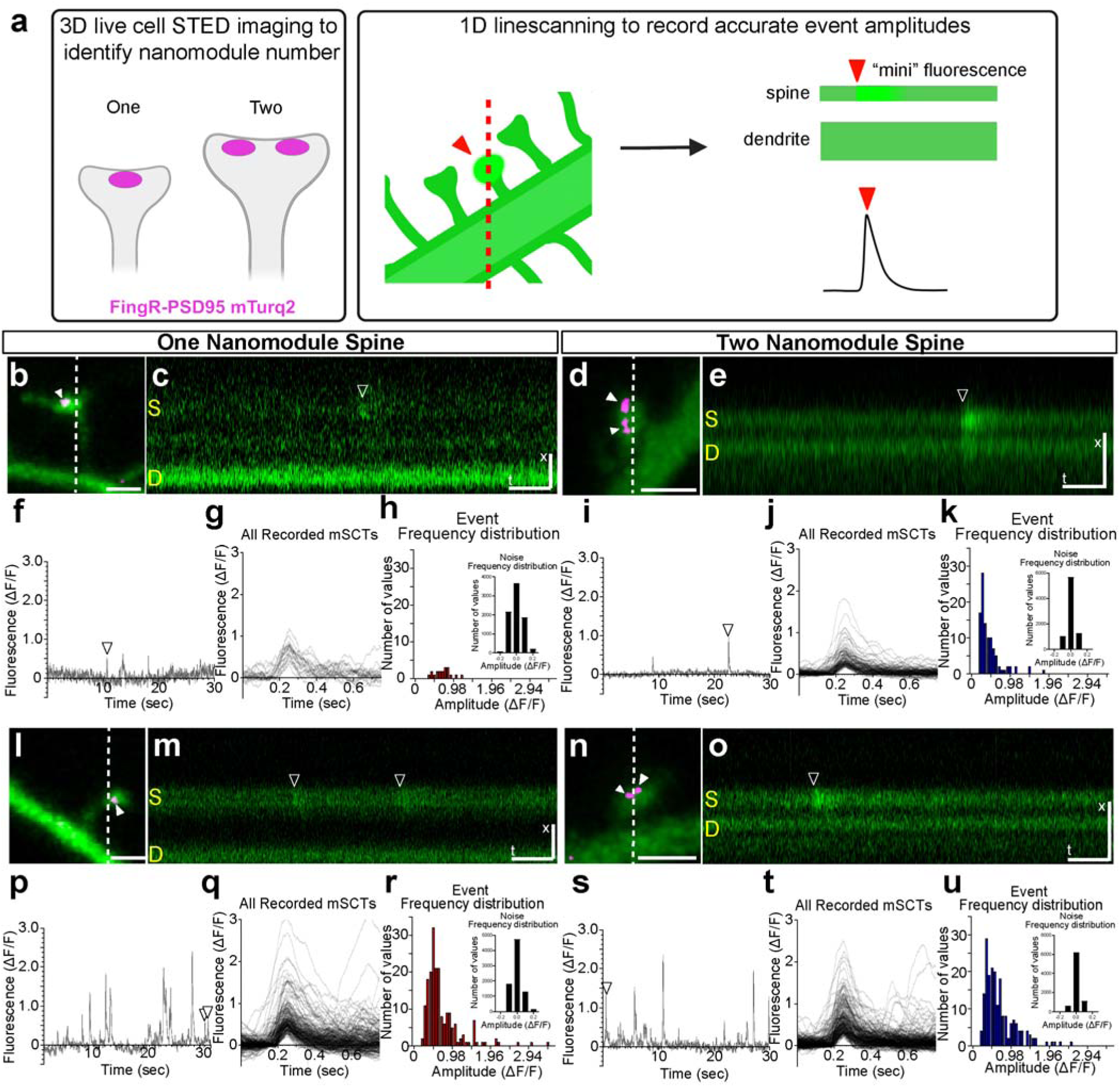
Quantification of mSCTs from individual dendritic spines. **(a)** Model depicting the two-step process of identifying nanomodule number in spines using STED nanoscopy, followed by drawing a line through selected spines and corresponding parent dendrite to record mSCT activity. One nanomodule **(b, l)** or two nanomodule **(d, n)** spines with GCaMP8f as a cell-fill (green) imaged with confocal, STED resolved PSD-95-FingR puncta (magenta, solid white arrow) and the dashed line indicating the position of the line scan used for high temporal resolution imaging. All scale bars for morphology are 1 μm. **(c, e, m, o)** Representative line-scans of the respective spines (b, d, l, n) with an mSCT shown (arrowhead). The spine and parent dendrite are marked (S = spine, D = dendrite). All scale bars for line scans are x = 1 μm and t = 200 ms. **(f, i, p, s)** Quantification of 30 seconds of recording from each respective spine with the corresponding event from representative line scans labeled (white arrow). **(g, j, p, t)** Quantification of every event recorded from each respective spine. **(h, k, r, u)** Amplitude histograms depicting all the recorded events from one nanomodule (red) and two nanomodule (blue) spines with 8000 points of baseline noise (black, inset). Models in (a) generated using BioRender.

The frequency of spontaneous events varied from spine to spine, but there was no difference in mean frequency between one and two nanomodule spines (Supplementary Fig. 4a). Therefore, we asked whether there were differences in events between spines sharing the same PSD-95 nano-organization. The amplitude of mSCTs in all spines, regardless of frequencies, exhibited positively skewed mSCT amplitude histograms, irrespective of the PSD-95 nanomodule number (lower frequency: Fig. 2g, j; higher frequency: Fig. 2r, u). Lower frequency spines with fewer events exhibited event amplitudes of similar sizes, resulting in a narrow distribution with a low standard deviation, consistent with single quantum events (Fig. 2h, k). Conversely, higher frequency spines with more events displayed event amplitudes that fell into multiple, distinct, evenly spaced ranges that increased progressively in size in a pattern reminiscent of multi-quantal release ^26–28^ (Fig. 2q, t). There was a weak negative correlation between mean mSCT frequency and mean amplitude in one nanomodule spines, while this negative correlation was not exhibited in two nanomodule spines (Supplementary Fig. 4b, c). Therefore, we hypothesized that spontaneous activity in both one and two nanomodule spines abides by quantal release characteristics. One limitation of this approach is that the spontaneous presynaptic response is being integrated using the activation of postsynaptic NMDAR, raising the possibility that the distribution of NMDAR mSCT amplitudes simply reflects the number of postsynaptic NMDARs being activated ^29^. Therefore, an approach was developed to measure the amount of glutamate released at synapses with identified nanoarchitecture.

### Vesicular glutamate release is quantal in both one and two nanomodule spines

To determine whether the quantal distribution of mSCTs depended on pre- or postsynaptic release characteristics, cortical neurons were transfected with the postsynaptically-targeted glutamate sensor, iGluSnFR3-SGZ^30^ (iGluSnFR3), along with PSD95-FingR to determine the relationship between presynaptic glutamate release and the nanoscale organization of PSD-95 at dendritic spines. One and two nanomodule spines were identified using STED (Fig. 3a, c, k, m). Miniature glutamate transients (mGluTs) were seen visually (Fig. 3b, d, l, n, Movies S9-S12) and quantifiably (Fig. 3e, h, o, r) from line scans. Line scan imaging of single spines resulted in a slightly higher rate of bleaching than GCaMP8f (4.6% mean reduction in raw fluorescence, Supplementary Fig. 3b), but small events were easily detectable throughout the imaging period. As seen with GCaMP8f, spines exhibited a range of event frequencies but there was no difference in mean frequency between one and two nanopuncta spines (Supplementary Fig. 4d). In spines with lower levels of activity, the distributions of mGluT amplitudes exhibited positive skew, irrespective of the number of PSD-95 nanomodules present. Notably, events from lower frequency spines were distributed within a narrow range (Fig. 3f, i), resulting in low standard deviation amplitude distributions (Fig. 3g, j) consistent with the release of a single quantum. In higher frequency spines, the mGluT amplitude distributions displayed a positive skew (Fig. 3q, t), consistent with multiple quantal events. Similar to mSCTs recorded using GCaMP8f, there was a weak negative correlation between mGluT frequency and amplitude in one nanomodule spines, which was reduced in two nanomodule spines (Supplementary Fig. 4e, f). Thus, findings in neurons transfected with iGluSnFR3 were consistent with results using GCaMP8f, suggesting that visualizing either miniature glutamate or NMDAR-dependent calcium events from single spines enables us to visualize spontaneous synaptic events at individual spines.

**Figure 3:**
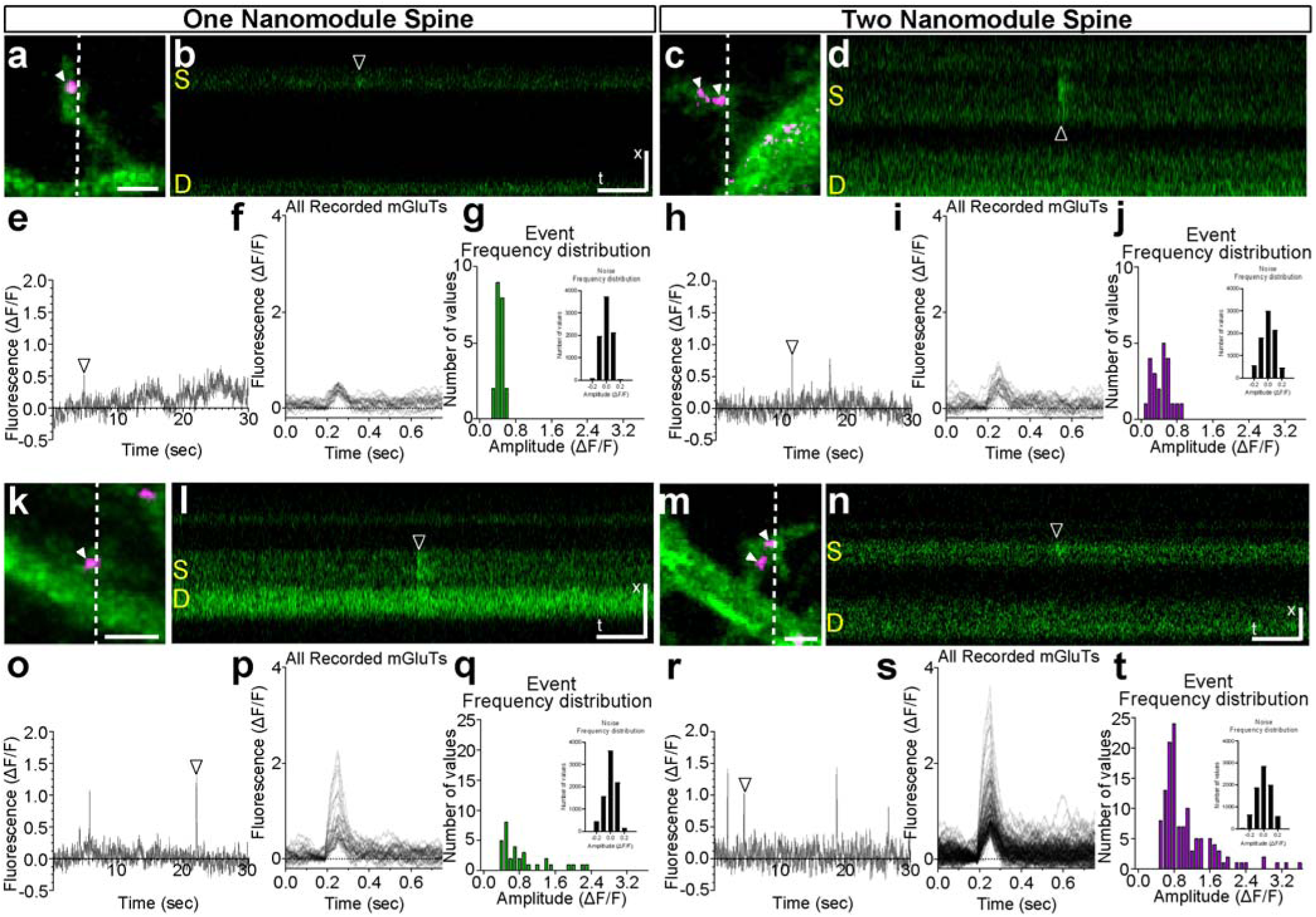
Quantification of mGluTs from individual dendritic spines. One nanomodule **(a, k)** or two nanomodule **(c, m)** spines with iGluSnFR3 as a cell-fill (green) imaged with confocal, a STED resolved PSD-95 FingR puncta (magenta, solid white arrow), and a dashed line indicating the position of the line scan used for high temporal resolution imaging. Scale bar is 1 μm. **(b, d, l, n)** Representative line-scans of the corresponding spines (a, c, k, m) with an mGluT shown (solid black arrow). The spine and parent dendrite are marked (S = spine, D = dendrite). Scale bar is x = 1 μm and t = 200 ms. **(e, h, o, r)** Quantification of 30 seconds of recording from each respective spine with the corresponding event from the representative line-scan labeled (white arrow). **(f, i, p, s)** Quantification of every event recorded from each respective spine. **(g, j, q, t)** Peak amplitude histogram depicting all the recorded events from the respective one nanomodule (green) or two nanomodule (magenta) spines with 8000 points of baseline noise (black, inset).

### One and two nanomodule spontaneous activity exhibit no kinetic differences

We determined the event kinetics of miniature events, including rise times, decay times, and FWHM in one and two nanomodule spines. We hypothesized that there would be no differences in miniature event kinetics between one and two nanomodule spines, consistent with a model where nanomodules represent unitary sites of synaptic transmission. To test whether event kinetics were independent of nanomodule number, the rise times, decay times, FWHM time courses were calculated. There was no difference in these parameters of mSCTs and mGluTs between one and two nanomodule spines (Supplementary Fig. 5c-e, h-j). Likely due to the buffering of intracellular calcium, which increases with respect to spine volume^16^, when mSCTs were visualized in two nanomodules spines, their apparent peak amplitudes were less than mSCT amplitudes in one nanomodule spines, (Supplementary Fig. 5a, b).The event amplitude of mGluTs was not different in one or two nanomodule spines (Supplementary Fig. 5f, g) due to iGluSnFR3 being expressed on the postsynaptic surface. All of the recorded mGluTs and mSCTs were scaled to the same size between nanomodule groups to further visualize differences between waveforms from one and two nanomodule spines. There was no difference in the waveforms of scaled events (Supplementary Fig. 6a-j), consistent with a model where the kinetics of mSCTs or mGluTs events do not depend on nanomodule number. This suggested nanoclusters of PSD-95 may reflect the number of independent release sites at a synapse.

### Quantal analysis of individual spines resolves quantal release characteristics

To test whether the nanoscale organization of spines relates to synaptic function, distributions of mSCT and mGluT peak amplitudes at single dendritic spines were examined. In spines with larger numbers of events, the distribution of event size appeared to segregate into discrete, evenly spaced amplitude bins. These data were consistent with the model that synaptic events represented the quantal fusion of synaptic vesicles^31^. However, single spines could not be imaged long enough to visualize a sufficient number of events for quantal analysis. Therefore, we sought an approach to increase the number of events recorded. Since the kinetics of single events were the same in all spines, events recorded from all single spines in neurons expressing either GCaMP8f or iGluSnFR3 were combined into a single mSCT or mGluT amplitude histogram, respectively. Consistent with previous reports that did not account for the nanoscale organization of spines^32^, combining all events – irrespective of nanomodule number – showed that the event data for both GCaMP8f and iGluSnFR3 events were well fit by both a skewed Gaussian distribution or a multiple term Poisson distribution (Fig. 4a, b). However, further separating spines by nanomodule number allowed for better visualization of quantal peaks in the combined amplitude histograms (Supplementary Fig. 8a-d). Despite this improvement, differences in baseline noise and quantal size between individual spines contributed to a smearing effect when amplitude histograms were aggregated. To address this, quantal analysis was performed to normalize these differences between spines and enhance visualization of quantal peaks.

**Figure 4:**
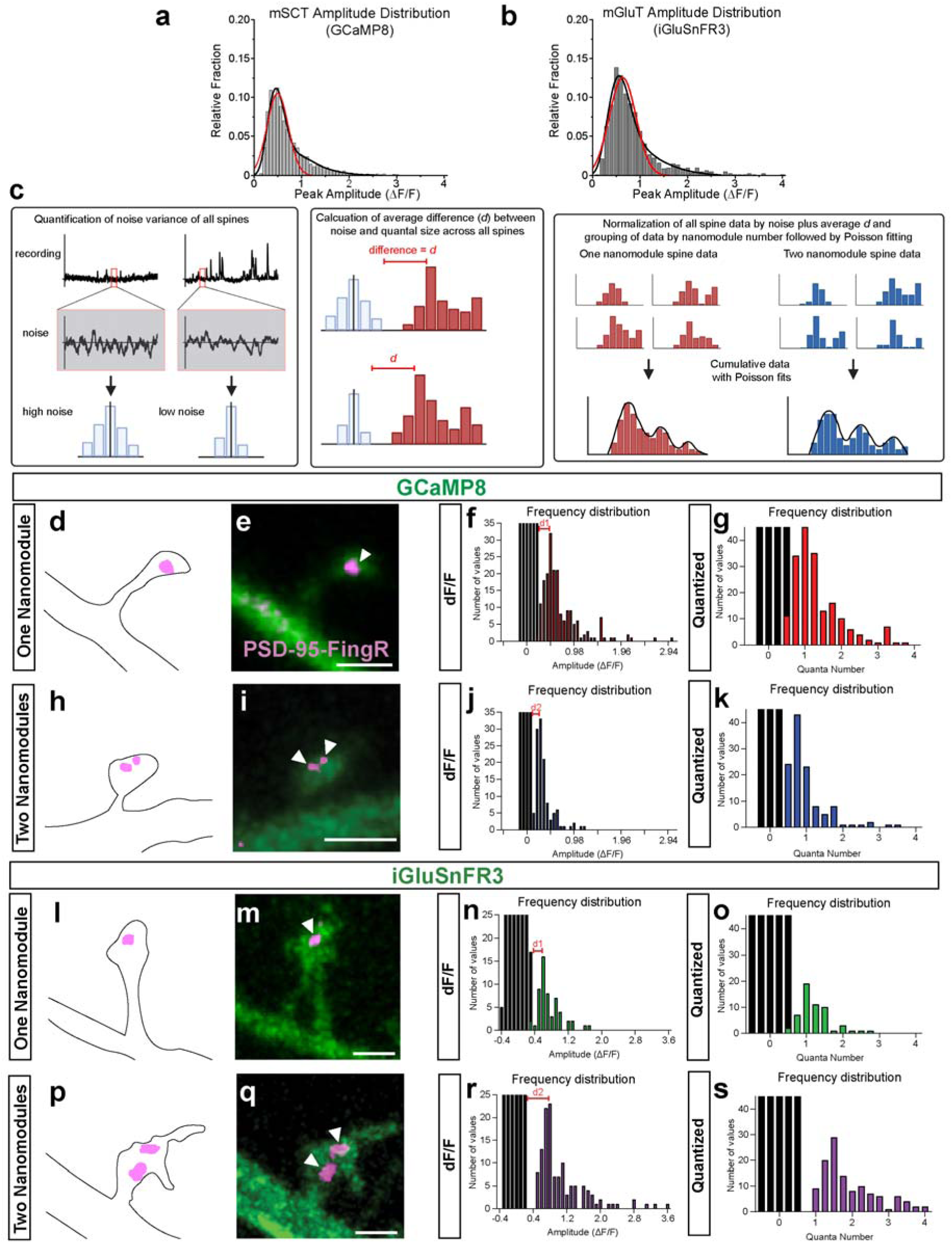
Noise characteristics of spines reveals non-Gaussian amplitude distribution. **(a)** Raw peak amplitude histogram from all spines recorded using GCaMP8f, with a skewed Gaussian distribution (red line) and a three quanta Poisson fit model (black line). N = 25 spines, 2164 events; Gaussian R^2^ = 0.889, AIC = −509.96, three quanta Poisson R^2^ = 0.976, AIC = −594.67. **(b)** Raw peak amplitude histogram from all spines recorded using iGluSnFR3-SGZ with a skewed Gaussian distribution (red line) and a three quanta Poisson fit model (black line). N = 23 spines, 799 events; Gaussian R^2^ = 0.934, AIC = −374.36, three quanta Poisson R^2^ = 0.977, AIC = −415.71. **(c)** Model depicting the process by which quantal analysis was performed, accounting for individual noise and quantal size variation of spines. **(d, e)** Examples of one nanomodule spines expressing GCaMP8f showing the PSD-95 puncta in magenta and spine as an outline (d) or in green (e). (**f**) Frequency distribution of event sizes from the spine (d, e) with recorded events (red) and noise (black) shown as histograms. **(g)** Quantized events from spine in **d-e**. Difference between noise and quantal size is also shown (red line, d_1_, d_2_). **(f, g)** Same as in **d**, **e** but for two nanomodule spines with recorded events (blue) and noise (black). **(h-k)** As in **d-g** but showing a spine with two PSD-95 puncta. (**l**, **m**) Examples of a one nanomodule spine as in **d, e** but expressing iGluSnFR3. (**n, o**) Histograms are of recorded iGluSnFR3 events (green) and noise (black) from the spine shown. **(p-s)** Same as in **l-o** but shows a two nanomodule spine. Models in **c** generated using BioRender. Scale bars = 1 µm.

Quantal analyses of each individual spine were performed as follows. The single quantal event size in each spine was determined by defining a quantum as the minimal detectable amplitude of the spontaneous events relative to the baseline noise^31^ (x = 0, Fig. 4c). The size of the first quantum was determined for each spine by averaging the absolute values between noise and the first peaks c^/^, where quantal size *m* is determined as the ΔF/F fluorescence F divided by the sum of highest amplitude of the noise per spine, N, and c^/^ (*m* = F/(N + c^/^), Fig. 4d-g, l-o, Supplementary Fig. 8). In the resulting quantized mSCT and mGluT event histograms, the first peak in each spine histogram was at or near 1 (Fig. 4h-k, p-s), indicating, as expected, that the most frequent event for each synapse was a single quantum. These data are consistent with a model where spontaneous excitatory synaptic transmission at single spines consists of miniature synaptic events of a similar minimum size, regardless of the number of nanopuncta of PSD-95.

### Nanoscale organization of spines reflects a functional difference in minisynaptic event properties

To determine whether spines with different numbers of nanomodules have functional differences, the mSCT and mGluT events from one and two nanomodules spines were placed into separate groups. The median mSCT and mGluT event size between one and two nanomodule was not significantly different, which is consistent with one quantum events being the most frequent event size (Fig. 5c, i). If nanomodules represent distinct release sites, spines with increasing numbers of nanomodules should have more events with a larger number of quanta^33,34^. Consistent with this model, in two nanomodule spines, both mSCT and mGluT event amplitude cumulative probability distributions demonstrated significantly increased numbers of larger events than in one nanomodule spines (Fig. 5d, j). These data suggest that increased nanomodule number increases the number of synaptic release sites, linking the nanoscale organization of synaptic proteins to synaptic transmission.

**Figure 5:**
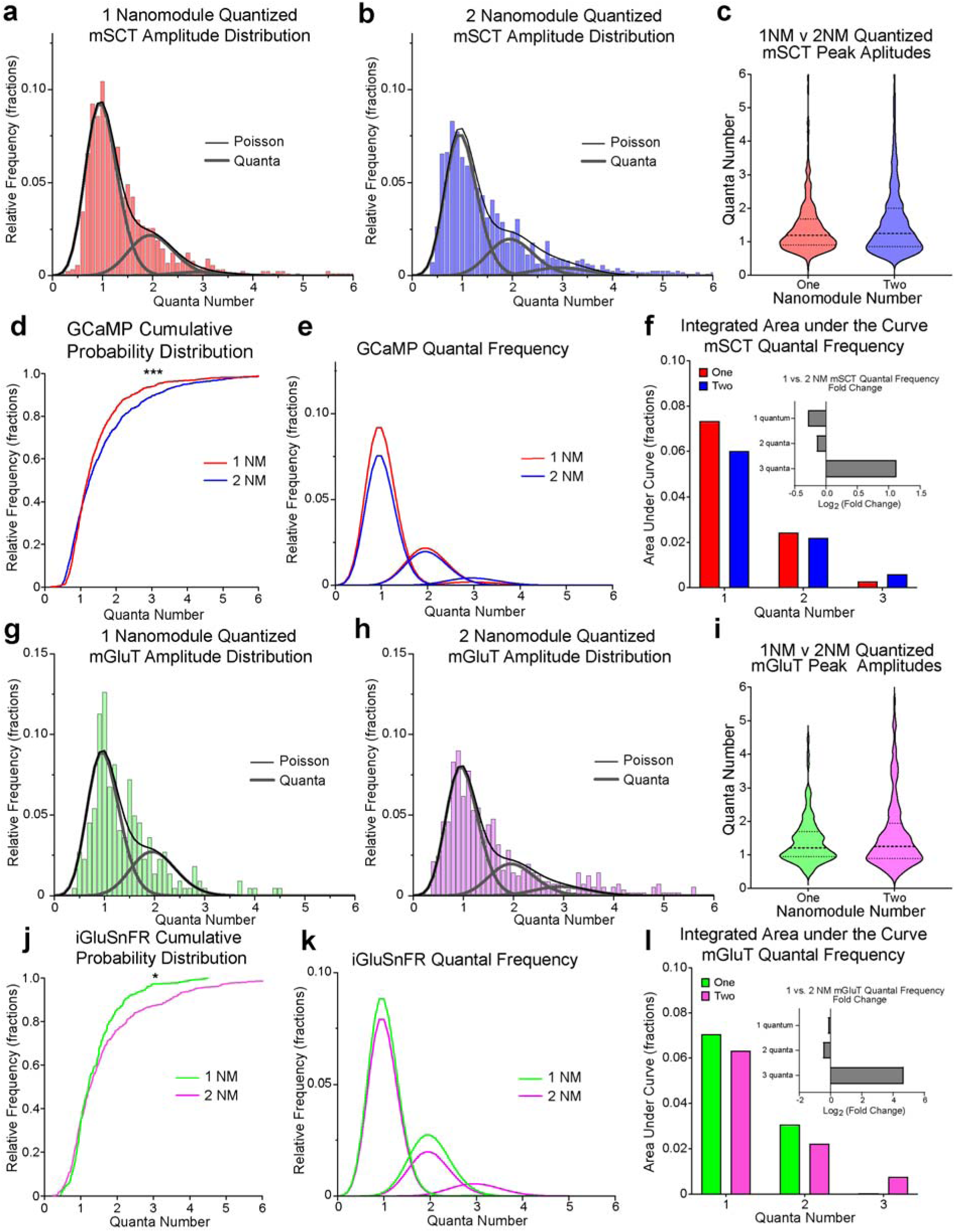
Comparing event amplitudes between populations of one and two nanomodule spines reveals quantal differences. **(a)** Quantized mSCT peak amplitude histogram from a population of one nanomodule GCaMP8f spines. Three quanta fit model is shown (black line, R^2^ = 0.962). Individual components of the function are shown in gray. N = 901 events across 11 spines. **(b)** Same as left (a), but for a population of two nanomodule spines (black line, three term Poisson R^2^ = 0.927). N = 1263 events across 14 spines. **(c)** Violin plots of quantized mSCT amplitudes from one nanomodule (red) and two nanomodule (blue) spines. Black dashed line represents quartiles. **(d)** Cumulative probability distributions of one nanomodule (red line) and two nanomodule (blue line) spines with respect to peak mSCT amplitudes (***p = 0.0004, by K-S test). **(e)** Individual quantal terms from Poisson distributions of one nanomodule (red line) and two nanomodule (blue line) GCaMP8f spines from above (a,b) overlaid upon each other. **(f)** The integrated area under the curve (AUC) calculated from each quantal term in (e) with the log_2_-fold change between the AUC of two nanomodule spines divided by one nanomodule spines (inset, fold change = (AUC two NM spines/AUC one NM spines)) **(g,h)** Quantized mGluT peak amplitude histograms from populations of one (g) and two (h) nanomodule iGluSnFR3 spines. Three quanta fit model is shown (black line, one nanomodule R^2^ = 0.848, two nanomodule R^2^ = 0.923) with individual components of the respective Poisson functions (gray). N = 222 events across 8 one nanomodule spines, N = 577 events across 15 two nanomodule spines. **(i)** Violin plots of quantized mGluT amplitudes from one nanomodule (green) and two nanomodule (magenta) spines. Black dashed line represents quartiles. **(j)** Cumulative probability distributions of one nanomodule (green line) and two nanomodule (magenta line) spines with respect to peak mSCT amplitudes (*p = 0.0308, by K-S test). **(k)** Individual quantal terms from the Poisson distributions of one nanomodule (green line) and two nanomodule (magenta line) iGluSnfR3 spines from above (f, g) overlaid upon each other. **(l)** Same integrated AUC as in (f) except based on mGluT data in (k).

To test whether these increases in event size reflected larger quantal events or more quantal release, Poisson distributions of mSCTs and mGluTs were determined for one or two nanomodule spines. Events from both one and two nanomodule spines for mSCTs and mGluTs were best fit by three quanta models (Fig. 5a, b, g, h). The three-term Poisson functions provided a better fit to the data than skewed Gaussian functions in all cases using Akaike Information Criterion (AIC) (Supplementary Fig. 9a, b, d, e). Additionally, the mSCT and mGluT amplitude histograms from all spines were well fit by three-term Poisson functions (Supplementary Fig. 10a, b). The Poisson distribution of two nanomodule spine mSCT events (Fig. 5e) and mGluT events (Fig. 5k) had fewer one- and two-quanta events and more three-quanta events than single PSD-95 nanopuncta spines. Spines with two nanomodules had a 2.2-fold increase in three quanta mSCTs and 25.3-fold increase in three quanta mGluTs when compared to one nanomodule spines (Fig. 5f, l). The data from two nanomodule spines were best fit by a three-quanta model, suggesting that the increased number of larger events was not due to the presence of larger (four+) quanta in two nanomodule spines. The increased number of three quanta events is consistent with the cooperativity of release, perhaps by reducing the energy barrier for vesicle fusion^26,35^. Regardless, these data support a model where both one and two nanomodule spines have multivesicular release, with an increasing number of nanopuncta leading to disproportionate increases in three quanta multivesicular release. These data support a model whereby synapses can modulate their pre- and postsynaptic signaling machinery to alter their strength.

## Discussion

The majority of spines contain a single nanomodule of synaptic proteins with <35% of spines containing two or more pre- and postsynaptic nanomodules^6^. The differences in nanoscale organization suggest that there may be functional differences between spines of different sizes^6,7^. Using a combined super-resolution and live-cell imaging approach, we find that the size of the smallest miniature calcium and glutamate events are the same, regardless of whether spines have one or two nanopuncta of PSD-95, suggesting that nanopuncta of PSD-95 mark independent synaptic release sites. The distributions of mSCT and mGluT amplitudes show similar quantal patterns, with both single and two nanomodule spines fit by three quanta Poisson distributions. Given that the fusion of single vesicles activates similar numbers of postsynaptic receptors, these data support the model that the precise nanoscale feature of spines are likely to be determinative for their function.

As spines become larger, they exhibit larger postsynaptic responses^1,2^. Previous work has suggested that these increases in synaptic strength are due largely to postsynaptic changes, for instance, in receptor number, subtype, and modulation^36–38^. Visualization of the mSCT and mGluT events in spines with known nanoarchitecture supports an additional mechanism for the increased size of postsynaptic responses in larger spines: coordination of multivesicular release. Spines with two nanopuncta of PSD-95 have significantly more three quanta events than spines with single nanopuncta, a striking increase in multiquanta events. Multivesicular release has been shown in central synapses both in the cortex and in the hippocampus^26–28,39,40^. Moreover, live-cell imaging of pre-and postsynaptic structures at the nanoscale reveals a coordinated increase in both pre-and postsynaptic nanopuncta following structural plasticity^6^. These data support the model that dendritic spine size is related to synaptic strength and the nanoscale organization of synaptic proteins. Taken together, these data suggest that each nanopuncta acts independently, and increases synaptic strength in larger spines may reflect a change in both presynaptic release machinery and postsynaptic receptors that is reflected in the nanoscale organization of the synapse.

Central nervous system synapses have long been considered to function differently from those in the periphery and to consist of only a single release site^41–45^. Our data suggest that nearly 35% of spines that contain more than one nanomodule of PSD-95 exhibit similarities to peripheral synapses and may contain multiple release sites. One challenge to the quantal neuromuscular junction model in the CNS has been the observation of a skewed distribution of miniature synaptic event amplitudes^32^. By segregating miniature synaptic responses by nanoarchitecture, the quantal characteristics of CNS synapses become apparent and suggest that nanostructural features are key functional units of spine synapses. In large spines with higher nanomodule number, there are more multivesicular events consistent with a model of a synapse that is functionally similar to peripheral synapses at the neuromuscular junction, where cortical nanocolumns resemble the multiple discrete active zones within lower motor neuron boutons^46^. Consistent with multivesicular release, EM analysis of CNS synapses has shown multiple endocytic pits ^47^. Our data provides a model of the central synapse; rather than a single release site that undergoes primarily postsynaptic modifications, we propose that the as the number of nanomodules of spine synapses increase there is an increase in coordinated vesicle release, making the classification of synapses based on their nanostructural features a necessary method to compare functional differences between spine synapses.

## Author contributions

Conceptualization: HJ and MBD; Methodology: HJ, SJT, SHY, REC, MBD; Investigation: HJ; Visualization: HJ, SJT; Funding acquisition: MBD; Supervision: MBD; Writing: HJ, SJT, MBD.

## Data and materials availability

All data are available in the main text or the supplementary materials. Analysis tools and imaging data available on request.

