## Supplemental Figures and Legends for "Nano-organization of synapses defines synaptic release properties at cortical neuron dendritic spines"

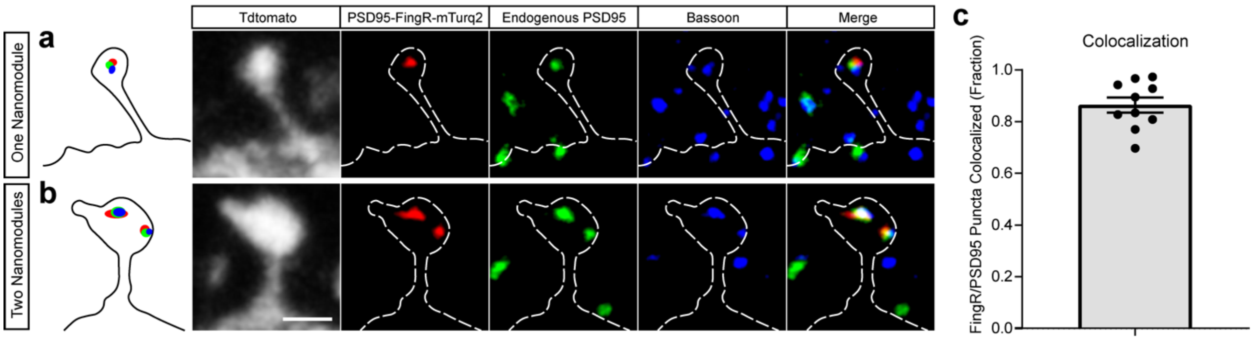


**Supplementary Figure 1. FingR-mTurq2 Colocalizes with Endogenous PSD95**

(**a**) Model of an exemplary one nanomodule spine (left) filled with TdTomato (gray) for morphology, with STED nanoscopy of immunostained PSD95-FingR-mTurq2 (red), endogenous PSD95 (green), and Bassoon (blue), and a merged image demonstrating colocalization of three puncta (right). White dashed outline represents morphology. **(b)** Model of an exemplary two nanomodule spine (left) with same immunostaining as shown above (a). Scale bar represents 1µm. **(c)** Fraction of FingR puncta which colocalized with endogenous PSD95 (mean = 86.4%, SEM = 2.9%). N = 10 cells, two regions per cells, a total of 983 PSD95-FingR-mTurq2 nanomodules were counted.

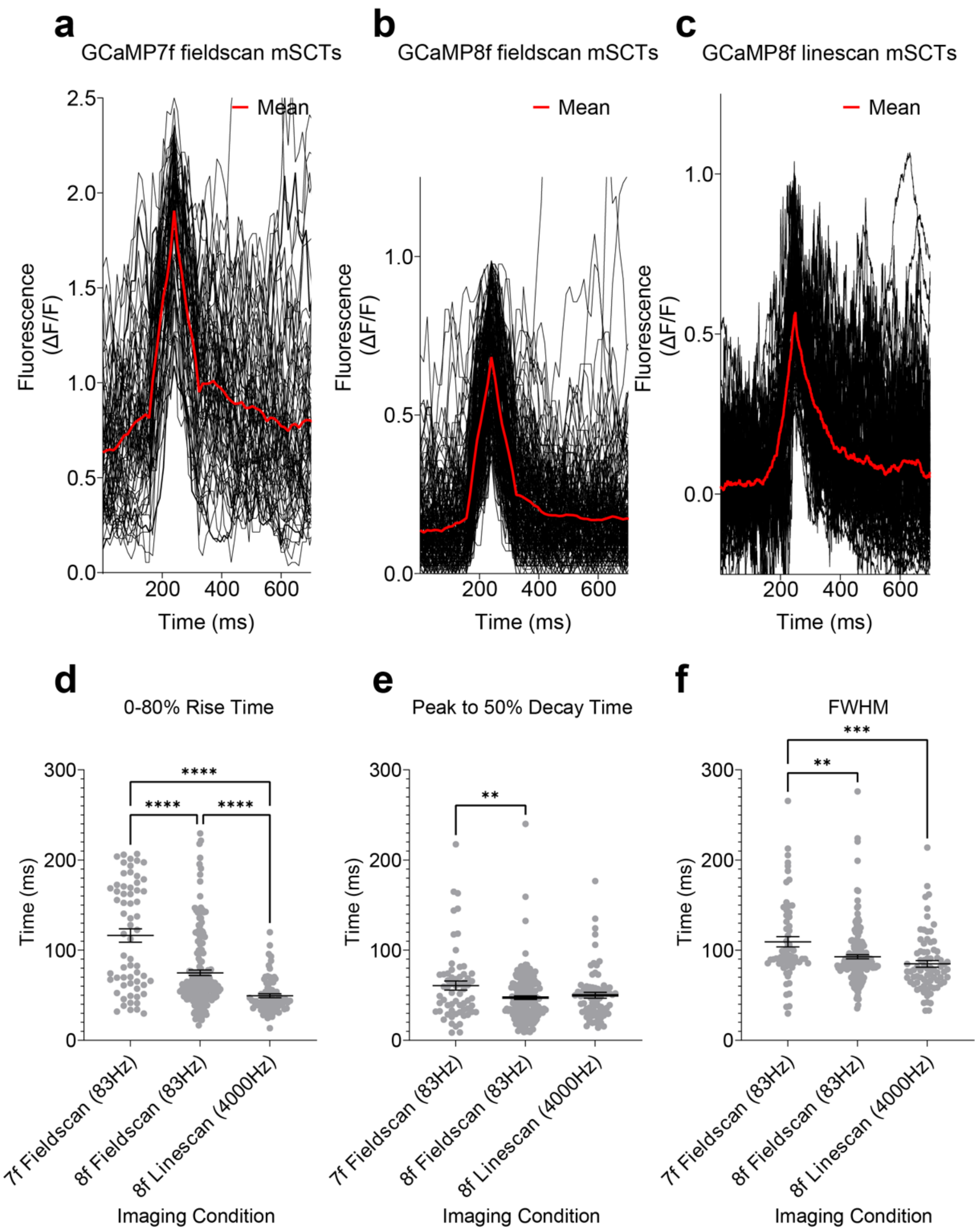


**Supplementary Figure 2. Kinetic Measurement of mSCTs using different Indicators and Acquisition Modes**

**(a)** Normalized ΔF/F fluorescence intensity of populations of small mSCTs. Individual events (black) recorded via 2-dimensional fieldscanning of dendritic spines expressing GCaMP7f displayed with the mean trace (red). Only the smallest events of the population of total events (not shown) were selected. Imaging was performed at 1 frame/12ms (~83Hz). N = 67 events. **(b)** Individual events (black) recorded via 2-dimensional fieldscanning of dendritic spines expressing GCaMP8f displayed with the mean trace (red). Only the smallest events of the population of total events (not shown) were selected. Imaging was performed at 1 frame/12ms (~83Hz). N = 184 events. **(c)** Individual events (black) recorded via 1-dimensional line scanning of dendritic spines expressing GCaMP8f displayed with the mean trace (red). Only the smallest events of the population of total events (not shown) were selected. Imaging was performed at 1 line/250µs (4000Hz). N = 78 events. **(d)** Quantification of the rise time from baseline to 80% of peak amplitude (GCaMP7f fieldscan mean = 116.2ms, SEM = 7.4ms; GCaMP8f fieldscan mean = 74.9ms, SEM = 3.1ms; GCaMP8f linescan mean = 49.5ms, SEM = 2.2ms)(****$p<0.0001$, One-way ANOVA, Tukey's posthoc). **(e)** Quantification of the decay time from the peak to 50% of peak amplitude (GCaMP7f fieldscan mean = 60.7ms, SEM = 5ms; GCaMP8f fieldscan mean = 47.4ms, SEM = 2ms; GCaMP8f linescan mean = 50ms, SEM = 3.2ms)(**$p = 0.0076$, One-way ANOVA, Tukey's posthoc). **(f)** Quantification of the full width at half maximum peak amplitude (GCaMP7f fieldscan mean = 109.3ms, SEM = 5.8ms; GCaMP8f fieldscan mean = 92.7ms, SEM = 2.3ms; GCaMP8f linescan mean = 84.8ms, SEM = 3.7ms)(**$p =0.0033$, ***$p = 0.0001$, One-way ANOVA, Tukey's posthoc).

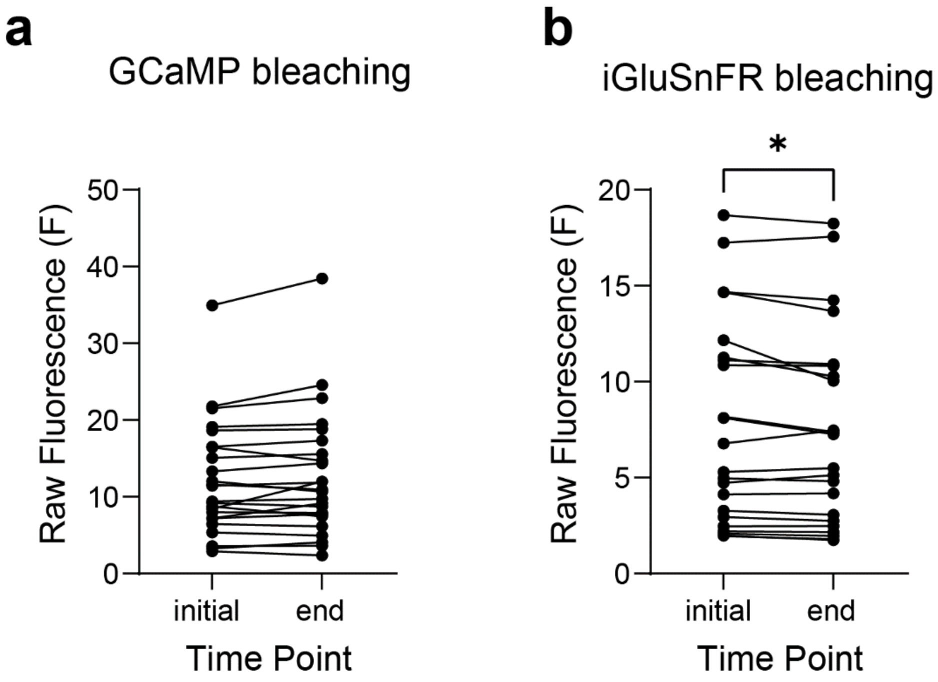


**Supplementary Figure 3. Photobleaching Ratio Across Imaging Runs**

**(a, b)** Raw fluorescence of ROIs drawn around parent dendrites of all spines that were recorded at an initial timepoint (first 5 seconds of imaging) and a final timepoint (last 5 seconds of imaging) for GCaMP8f (a) and iGluSnFR3-SGZ (b) (*p = 0.035, paired t-test).

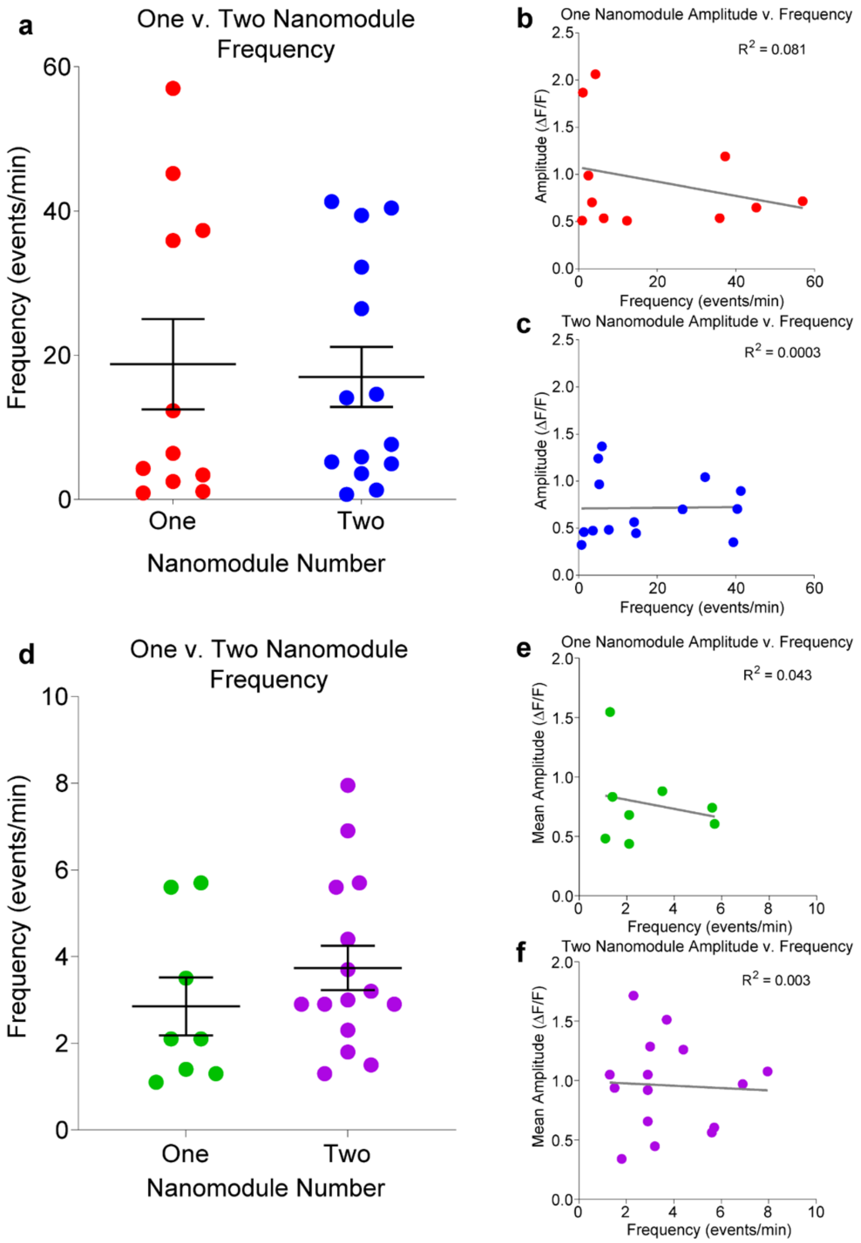


**Supplementary Figure 4. Frequency Differences of mSCTs and mGluTs Recorded from One and Two Nanomodule Spines**

**(a)** Comparison of mSCT frequency between one and two nanomodule spines. Mean = 18.8 events/min, SEM = 6.3 events/min, N =11 one nanomodule spines; mean = 17 events/min, SEM = 4.2 events/min, N = 14 two nanomodule spines. **(b)** Scatterplot comparing the ΔF/F normalized mean mSCT amplitude against the frequency for each one nanomodule spine. **(c)** Same as above (b) except for two nanomodule spines. **(d)** Comparison of mGluT frequency between one and two nanomodule spines. Mean = 2.9 events/min, SEM = 0.67 events/min, N = 8 one nanomodule spines; mean = 3.7 events/min, SEM = 0.51 events/min, N = 15 two nanomodule spines. **(e, f)** Scatterplots as shown above (b, c) comparing the ΔF/F normalized mean mGluT amplitude against the frequency for one and two nanomodule spines.

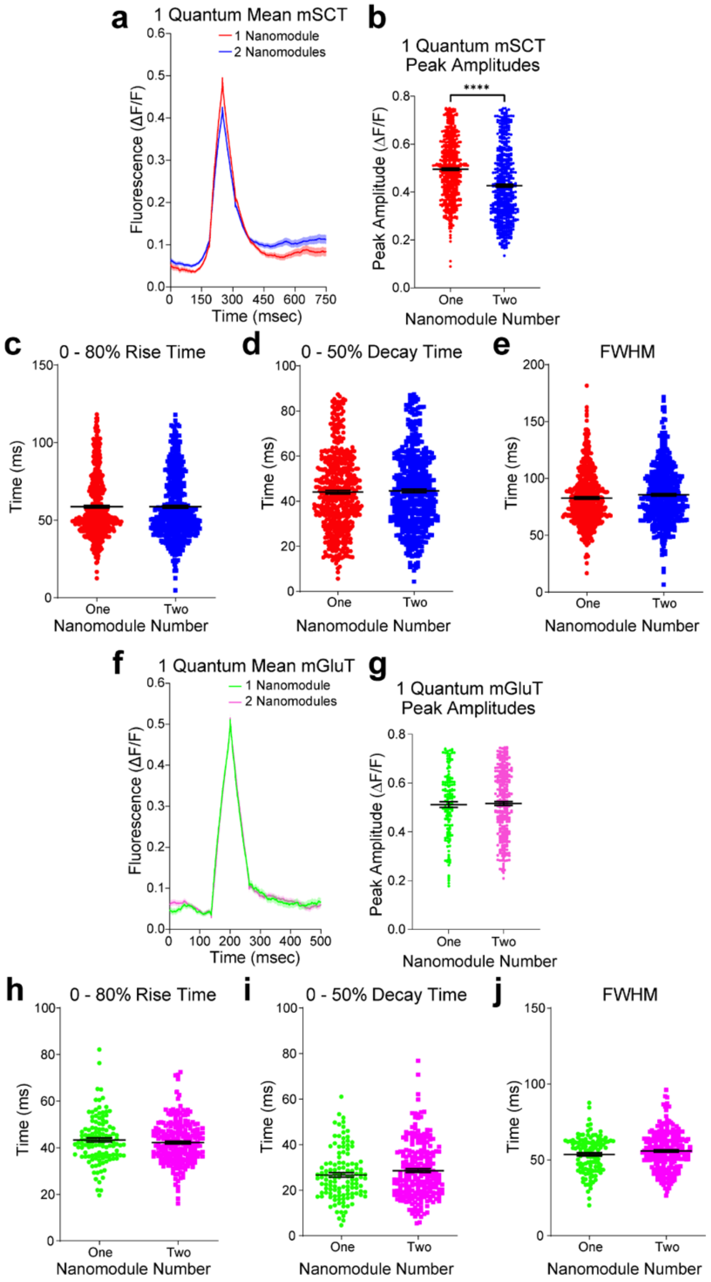


**Supplementary Figure 5. Kinetic Measurements of One Quantum mSCTs and mGluTs**

**(a)** Quantification of the mean waveform one quantum mSCT from one (red line) and two (blue line) nanomodule spines. N = 561 one nanomodule events, N = 558 two nanomodule events. **(b)** Peak mSCT amplitudes from the events averaged on the left (a). One nanomodule mean = 0.502, SEM = 0.006; two nanomodule mean = 0.433, SEM = 0.007; ****p < 0.0001, unpaired t-test. **(c)** Quantification of the one quantum mSCT rise time from baseline to 80% of peak amplitude. One nanomodule mean = 58.6ms, SEM = 0.9ms; two nanomodule mean = 58.6ms, SEM = 1.0ms. **(d)** Quantification of the decay time of one quantum mSCTs from the peak to 50% of peak amplitude. One nanomodule mean = 44ms, SEM = 0.8ms; two nanomodule mean = 44.6ms, SEM = 0.8ms. **(e)** Quantification of the full width at half maximum peak amplitude of one and two nanomodule mSCTs. One nanomodule mean = 82.3ms, SEM = 1ms; two nanomodule mean = 85.6ms, SEM = 1.1ms. **(f)** Quantification of the mean waveform one quantum mGluT from one (green line) and two (magenta line) nanomodule spines. N = 146 events from one nanomodule spines, N = 284 events from two nanomodule spines. **(g)** Peak mGluT amplitudes from the events averaged on the left (f). One nanomodule: mean = 0.512, SEM = 0.012; two nanomodule: mean = 0.516, SEM = 0.008. **(h)** Quantification of the one quantum mGluT rise time from baseline to 80% of peak amplitude. One nanomodule mean = 43.4ms, SEM = 0.9ms; two nanomodule mean = 42.3ms, SEM = 0.6ms. **(i)** Quantification of the decay time of one quantum mGluTs from the peak to 50% of the peak amplitude. One nanomodule mean = 26.7ms, SEM = 1ms; two nanomodule mean = 28.6ms, SEM = 0.8ms. **(j)** Quantification of the full width at half maximum peak amplitude of one and two nanomodule mGluTs. One nanomodule mean = 53.6ms, SEM = 1.1ms; two nanomodule mean = 55.9ms, SEM = 0.8ms.

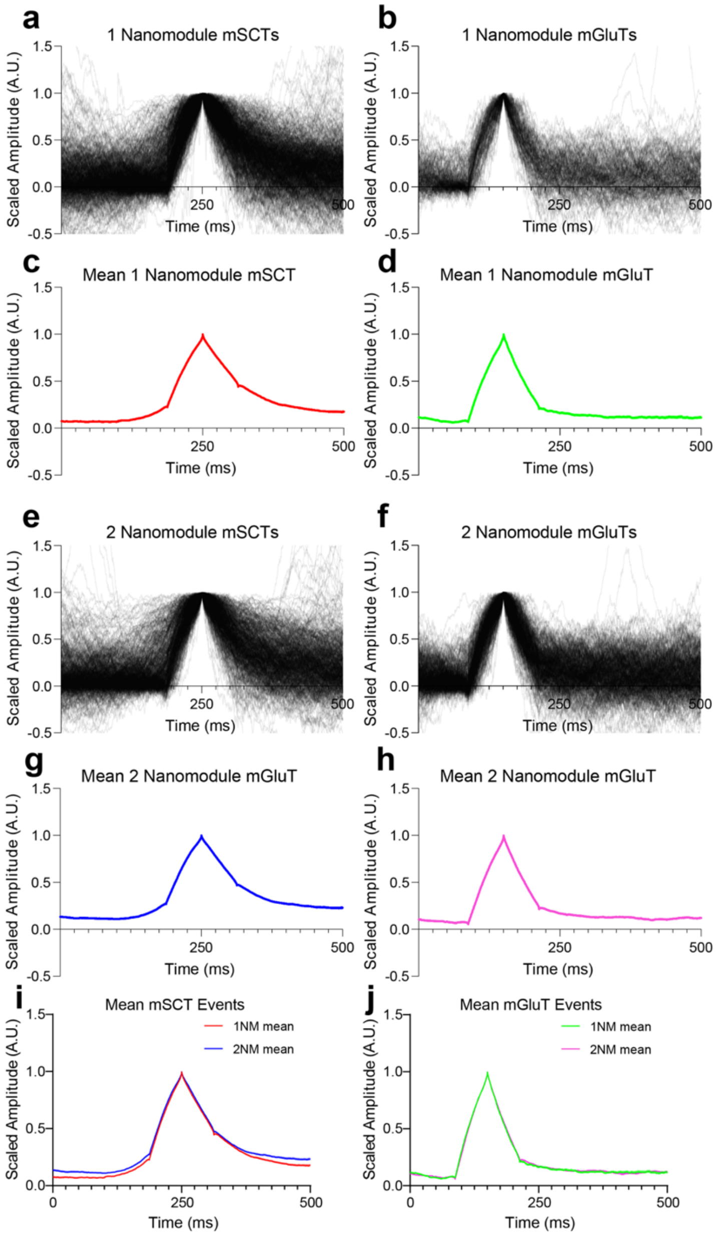


**Supplementary Figure 6. One and Two Nanomodule mGluTs Scaled to Equal Amplitudes**

(**a**) The mSCTs recorded from one nanomodule spines which were scaled to the same peak amplitude. N = 806 events. **(b)** The mGluTs recorded from one nanomodule spines which were scaled to the same peak amplitude. N = 222 events. **(c)** Averaged waveform from the events in (a). **(d)** Averaged waveform from the events in (b) **(e)** same as in (a) except for two nanomodule spines. N = 725 events. **(f)** Same as in (b) except for two nanomodule spines. N = 559 events. **(g)** same as in (c) except for two nanomodule spines. **(h)** Same as in (d) except for two nanomodule spines. **(i)** Overlay of mean mSCT traces in (c) and (g) from one (red) and two (blue) nanomodule spines. **(j)** Overlay of mean mGluT traces in (d) and (h) from one (green) and two (magenta) nanomodule spines.

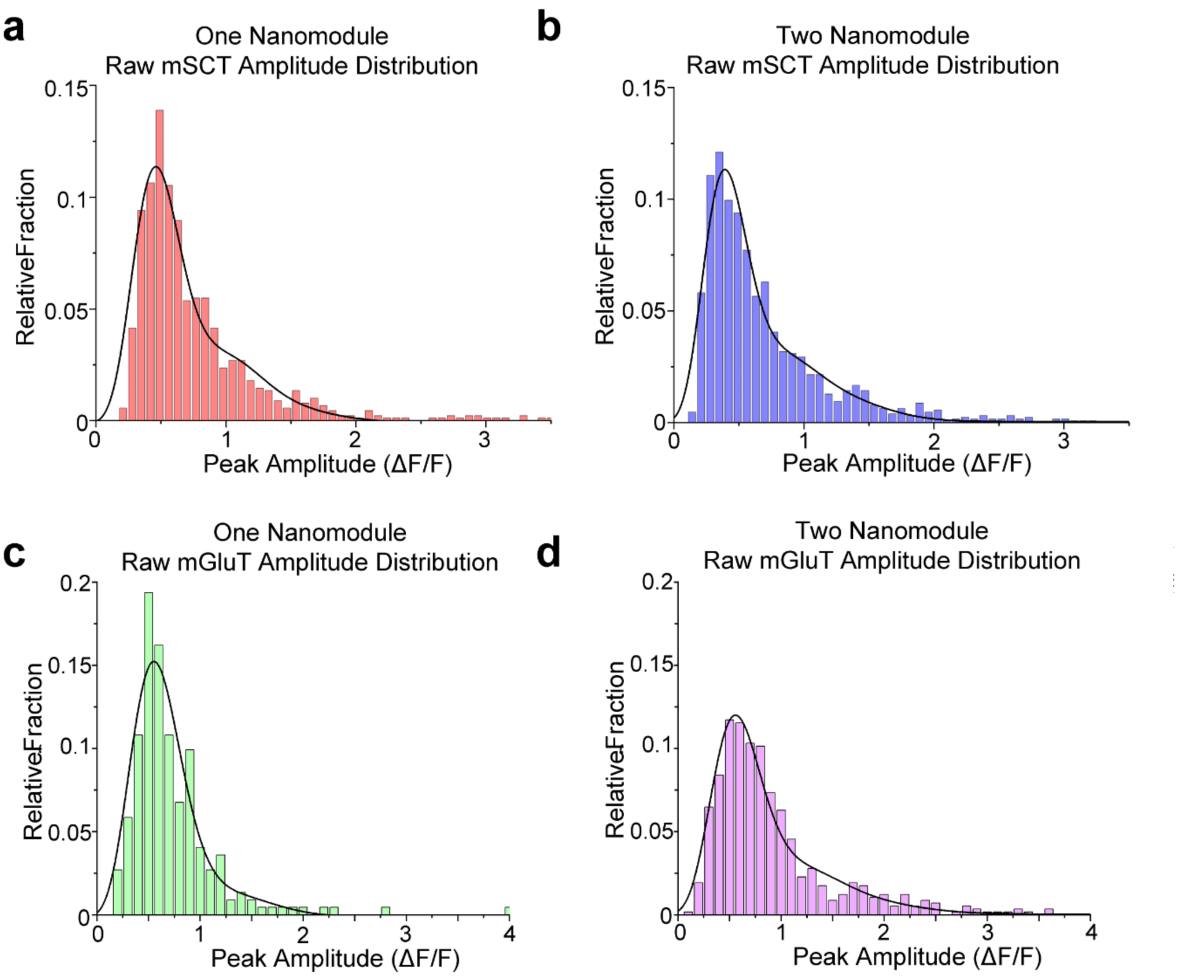


**Supplementary Figure 7. Raw Amplitude Histograms of Populations of One and Two Nanomodule Spines Using GCaMP8f and iGluSnFR3**

**(a, b)** Raw peak amplitude histograms from mSCTs from all one nanomodule (a) and two nanomodule. (b) spines, with 3 quanta Poisson fit models (black line). One nanomodule 3 quanta Poisson $R^2 = 0.932$; two nanomodule 3 quanta Poisson $R^2 = 0.957$ **(c, d)** Raw peak amplitude histograms from mGluTs from all one nanomodule (c) and two nanomodule (d) spines, with 3 quanta Poisson fit models (black line). One nanomodule 3 quanta Poisson $R^2 = 0.923$; two nanomodule 3 quanta Poisson $R^2 = 0.964$.

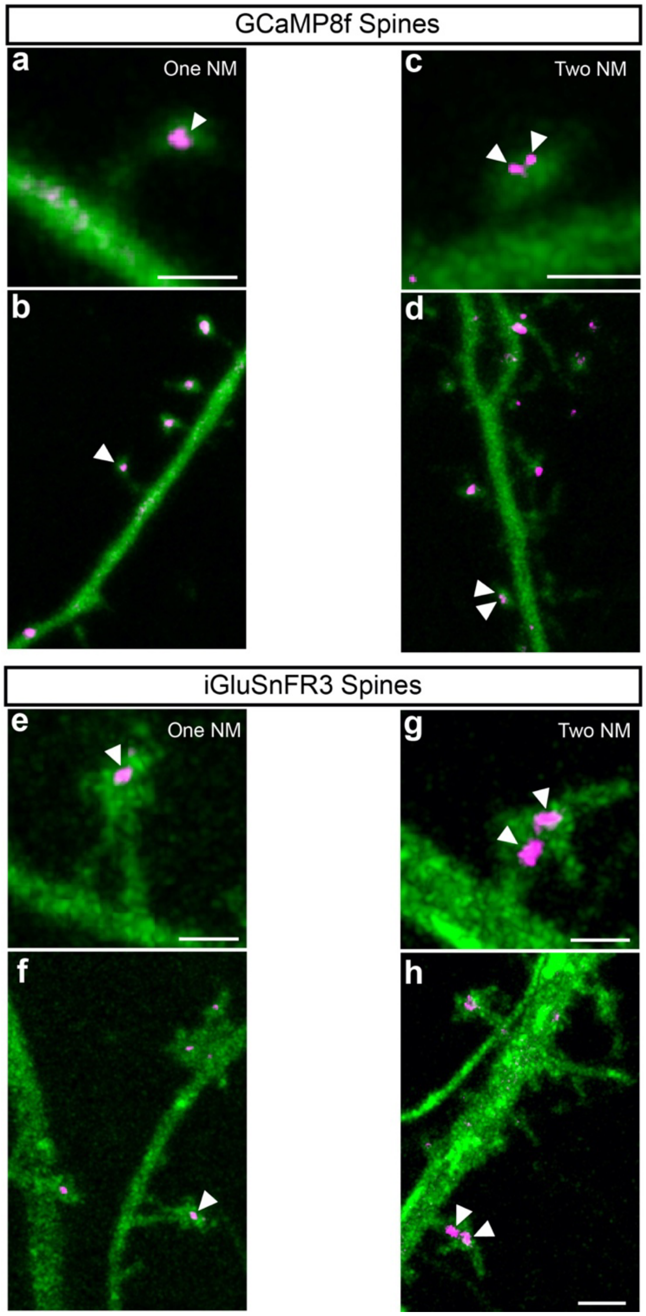


**Supplementary Figure 8. STED nano-organization of PSD-95 in spines low magnification images of dendrites from Figure 4**

All spines correspond to spines in figure 4. **(a, b)** One nanomodule GCaMP8f expressing spine corresponding to histogram data from **Fig 4d-g** with nano-organization shown (white arrows) **(c, d). c.** Shows image as in **Fig 4d** for reference. **d** Shows lower magnification image of dendritic region with the imaged spine. **(c,d)** As in **a,b** but the two nanomodule GCaMP8f expressing spine corresponding to histogram data from **Fig 4h-k** shown as in **a, b. (e,f)** As in a, b, but the mne nanomodule iGluSnFR3 expressing spine shown in **Fig 4i-o. (k, l)** As in a, b, but the two nanomodule iGluSnFR3 expressing spine corresponding to histogram data from **Fig4p-s** is shown. All scale bars = 1 µm.

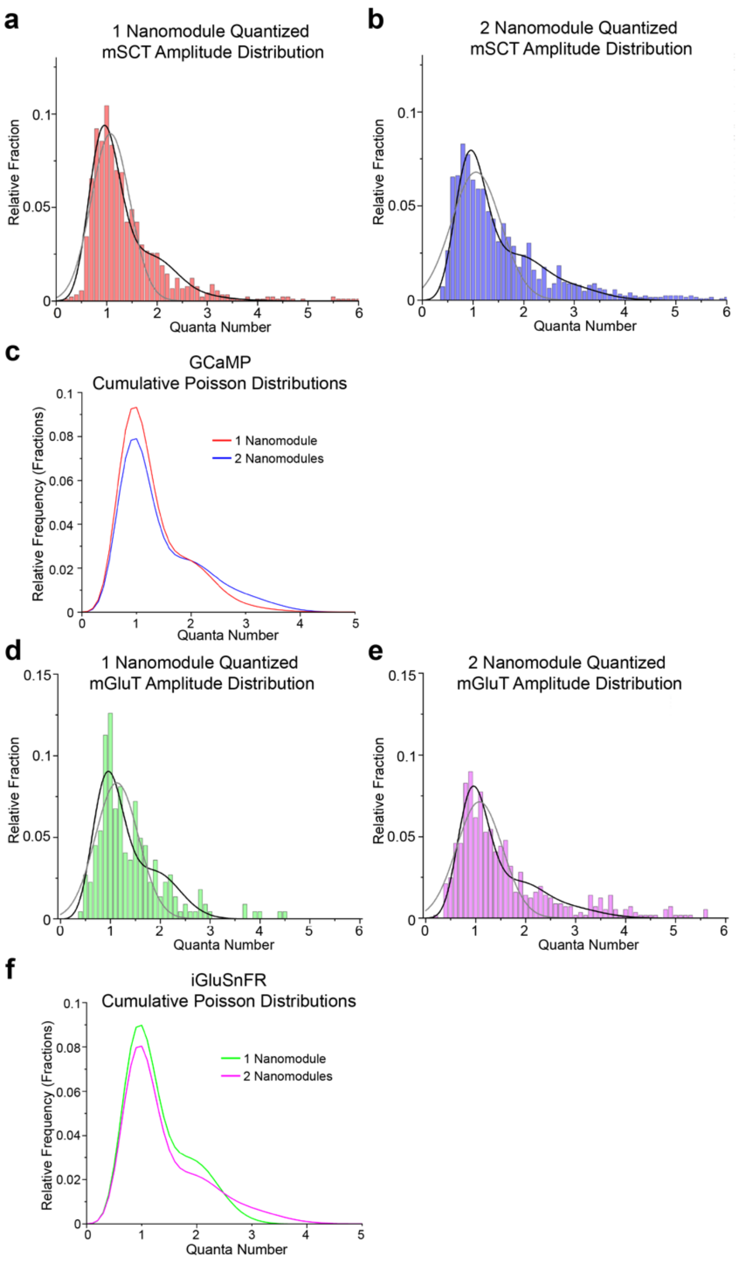


**Supplementary Figure 9. Cumulative Poisson Fits of Quantized Data from One and Two Nanomodule Spines**

(**a**) Quantized mSCT peak amplitude histogram from a population of one nanomodule spines. Three quanta fit model is shown (black line, $R^2 = 0.962$, AIC = -861.71). This was compared to the best fit Gaussian (gray line, $R^2 = 0.896$, AIC = -781.39). N = 901 events across 11 spines. **(b)** Same as left (a), but for a population of two nanomodule spines. Gaussian $R^2 = 0.8$, AIC = -557.36; three term Poisson $R^2 = 0.927$, AIC = -617.12). N = 1263 events across 14 spines. **(c)** Cumulative poisson distributions of quantized peak amplitudes from one nanomodule (red line) and two nanomodule (blue line) spines from above (a,b) overlaid upon each other. **(d,e)** Quantized mGluT peak amplitude histograms from populations of one (d) and two

(e) nanomodule spines. Three quanta fit model is shown (black line, one nanomodule $R^2$ = 0.848, AIC = -545.16, two nanomodule $R^2$ = 0.923, AIC = -611.32), and compared to the Gaussian fit (gray line, one nanomodule $R^2$ = 0.785, AIC = -525.24, two nanomodule $R^2$ = 0.838, AIC = -569.71). N = 222 events across 8 one nanomodule spines, N = 577 events across 15 two nanomodule spines. **(f)** Cumulative poisson distributions of quantized peak amplitudes from one nanomodule (green line) and two nanomodule (magenta line) spines from above (d, e) overlaid upon each other.

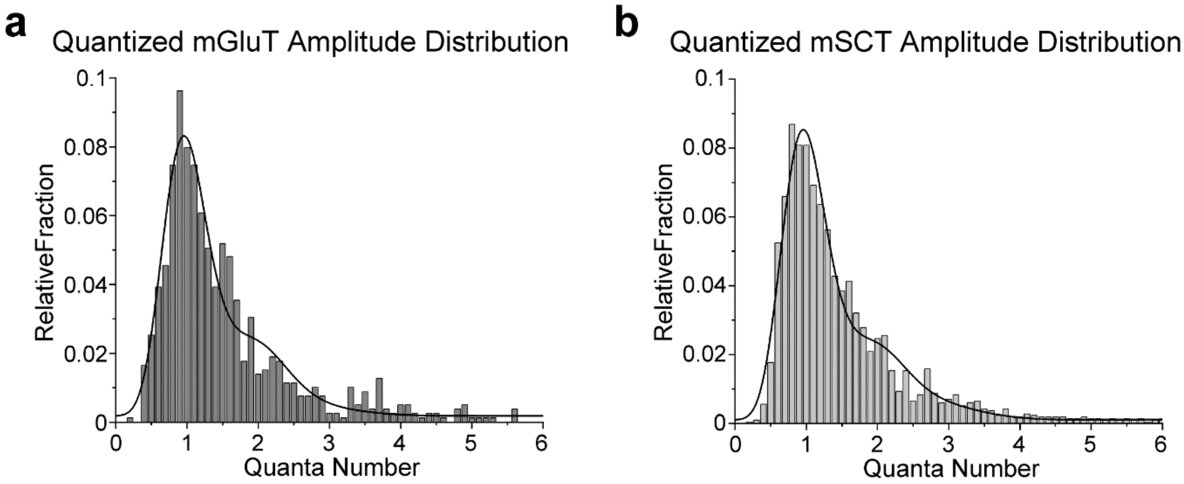


**Supplementary Figure 10. Quantized Peak Amplitude Histograms from Dendritic Spines not Separated by Nano-Organization**

**(a)** Quantized peak amplitude histogram of mGluTs from all spines with three quanta Poisson fit model (black line). N = 23 spines, 799 events, $R^2 = 0.944$. **(b)** Quantized peak amplitude histogram of of mSCTs from all spines with three quanta Poisson fit model (black line). N = 25 spines, 2164 events, $R^2 = 0.97$.

### Supplementary Movies

**Movie S1 – Video GCaMP8F signals in spine in Figure 1c.** Spine contains one nanopuncta of PSD-95.

**Movie S2 – Video GCaMP8F signals in spine in Figure 1d.** Spine contains two nanopuncta of PSD-95.

**Movie S3 – Video GCaMP8F signals in spine in Figure 1g.** Movie showing the same spine as in 1c following addition of APV.

**Movie S4 – Video GCaMP8F signals in spine in Figure 1h.** Movie showing the same spine as in 1d following addition of APV.

**Movie S5 – Line scans of GCaMP8f used for analysis shown in 2f containing example image shown in 2c.** Arrow marks event shown in 2c.

**Movie S6 – Line scans of GCaMP8f used for analysis shown in 2i containing example image shown in 2e.** Arrow marks event shown in 2e.

**Movie S7 – Line scans of GCaMP8f used for analysis shown in 2p containing example image shown in 2m.** Arrow marks event shown in 2m.

**Movie S8 – Line scans of GCaMP8f used for analysis shown in 2s containing example image shown in 2o.** Arrow marks event shown in 2o.

**Movie S9 – Line scans of GluSnfr used for analysis shown in 3e containing example image shown in 3b.** Arrow marks event shown in 3b.

**Movie S10 – Line scans of GluSnfr used for analysis shown in 3h containing example image shown in 3d.** Arrow marks event shown in 3d.

**Movie S11 – Line scans of GluSnfr used for analysis shown in 3o containing example image shown in 3l.** Arrow marks event shown in 3l.

**Movie S12 – Line scans of GluSnfr used for analysis shown in 3r containing example image shown in 3n.** Arrow marks event shown in 3n.
