## Supplementary material for "Nano-organization of synapses defines synaptic release properties at cortical neuron dendritic spines": Methods

**Animals**

All animal studies were approved by the Institutional Animal Care and Use Committee of Thomas Jefferson University (01286-1 and 012889-1). Long Evans embryonic day 17-18 (E17-18) rat embryos were harvested from timed pregnant rats (Charles River Laboratories, Wilmington, MA) and used to prepare primary cortical neuron cultures as described below.

**Primary cortical neuron culture preparation**

Primary cortical neurons were prepared as previously described (Hruska et al., 2022a; Hruska et al., 2018) from E17-18 rat cortexes. Neurons were cultured in Neurobasal Medium (cat #: 21103049, Thermo Fisher Scientific) supplemented with 1x B27 (cat #: 17504044, Thermo Fisher Scientific), 1x GlutaMAX (cat #: 35050061, Thermo Fisher Scientific) and penicillin-streptomycin (cat #: 15140-122, 10,000 U/mL, Thermo Fisher Scientific). Neurons were plated on poly-D-lysine (cat #: 354210, Corning, Corning, NY) and laminin (cat #: CB40232, Corning) coated 35 mm glass bottom dishes with a 13 mm well with #1.5 cover glass (cat #: GBD00002-200) at a density of 200,000 neurons/dish. Neurons were maintained in a humidified 37°C incubator with 5% $CO_2$ and transfected as described below.

**Plasmids and Plasmid Construction**

pSyn-Tdtomato (a generous gift from Peter Scheiffele, University of Basel, Biozentrum) was used to visualize neuronal morphology as previously described (Hruska M et al 2015, Hruska M et al 2018). In order to visualize endogenous PSD-95, we generated a pCAG-PSD95-FingR-mTurq2 construct. This was created using the pCAG-PSD95-FingR-eGFP construct (Addgene plasmid no. 46295) and swapping out eGFP for mTurq2. Additionally, the linker region was

**Commented [ST1]:** You need to enter these citations into EndNote extended by 19 amino acids upstream of the mTurq2. The calcium indicators pGP-AAV-synjGCaMP8f-WPRE (Addgene plasmid no. 162376) or pSyn-GCaMP7f were used for calcium imaging experiments. GATEWAY cloning (Thermo Fisher Scientific) was used to generate pSyn-GCaMP7f using a pCMV-GCaMP7f (generously provided by the lab of Tim Mosca, Thomas Jefferson University, Philadelphia, PA), as a donor vector and pSyn (previously generated in the lab) as a destination vector. The glutamate sensor pAAV-hSyn-iGluSnFR3v857-SGZ (Addgene plasmid no. 178330) was used for glutamate imaging experiments.

**Transfection of Primary Cortical Neurons**

Transfections were performed at DIV17-19 after a majority of spine formation had already occurred. 100 μL of Neurobasal media without supplements was added to two tubes corresponding to each dish being transfected. In one tube, DNA constructs were added. In the other corresponding tube, 0.5 μL of Lipofectamine 2000 (cat #: 11668027, Thermo Fisher Scientific) was added. After 5 minutes of incubation at room temperature, the DNA was added slowly dropwise to the tube containing Lipofectamine. Air was then gently bubbled through the mixture using a micropipette. The mixture was incubated for a further 15 minutes at room temperature. During incubation, the cells were removed from the cell incubator and the conditioned media was removed and saved. Then 600 μL of warm Neurobasal without supplements was immediately added to the cells. The conditioned media was stored at 37° C for later use. 200μL of the DNA/Lipofectamine mixture was added dropwise to the cells. The cells were then placed back in the incubator for 1.5-2 hours. After incubation, the warm conditioned media was sterilized through a 0.22 µm syringe filter. The DNA/Lipofectamine mixture was aspirated away and the filtered conditioned media was added back to the cells.

For the FingR localization experiments, 60 ng of pSyn-Tdtomato, as described in Hruska M et al 2018 and Hruska M et al 2015, and 50 ng of pCAG-PSD95-FingR-mTurq2 were used per dish. For the calcium imaging experiments, 150 ng of pGP-AAV-synjGCaMP8f-WPRE or pSyn-GCaMP7f and 50 ng of pCAG-PSD95-FingR-mTurq2 were used per dish. For the glutamate imaging experiments 300 ng of pAAV-hSyn-iGluSnFR3v857-SGZ and 50 ng of pCAG-PSD95-FingR-mTurq2 were used per dish.

**STED Imaging**

Live cell STED experiments were performed using a Leica TCS SP8 gated STED (GSTED) 3x super-resolution system (Leica Microsystems, Mannheim, Germany) equipped with a tunable pulsed white light laser, continuous wavelength (CW) 592 nm and 660 nm depletion lines and a pulsed 775 nm depletion line. Live cell confocal images of GCaMP7f, GCaMP8f, or iGluSnFR3 were acquired as z-stacks (190 nm slice thickness) with a 100× oil immersion objective (Leica Microsystems) using a resonant scanner (8,000 Hz), and gated HyD detectors (set at 200-350% gain) were set between 0.4 and 6 ns. Live cell STED images of FingR-PSD95-mTurq2 were taken using the same specifications, but without gating the detectors due to mTurq2 excitation using a CW 442 nm diode. A pixel format of 1024 x 1024 pixels and 4-5x zoom was used to achieve a pixel size of 25 nm. The 592 nm CW STED line was used to deplete mTurq2 (at 15-35% power) to achieve STED resolution.

For three-color gated STED images of fixed cultured neurons imaged using the Leica SP8 3 × GSTED, mTurq2 was labeled with Alexa-488, Bassoon was labeled with Alexa-594, and PSD-95 was labeled with Atto-647N. Resonance scanning (8,000 Hz), gated HyD detectors (set at 100–250%) and a 100× oil immersion objective (Leica Microsystems) with a pixel format of 1024x1024 pixels and a 4-5x zoom to obtain desired pixel size (25 nm) was used to acquire z-stacks of 190 nm slice thickness. Gated HyD detectors were set at 0.4-6 ns, and mTurq2 was excited with the 488 nm line (1-3% power), and the CW 592 nm line (30-45% power) was used to achieve STED resolution. Gated HyD detectors set at 0.3-6 ns were used to acquire Bassoon which was excited with the 594 nm laser line (8-15% power). The same detector settings were used to acquire PSD-95, which was excited with the 647 nm laser line (15-30% power). In order to achieve STED resolution for Bassoon and PSD-95, the pulsed 775 nm depletion line (30-40% for Bassoon and 20-30% for PSD-95) was used to generate a resolution of ~50 nm.
For *z* depletion, 10% of the depletion line power was redirected to the *z* donut to achieve an image *z*-resolved at ~250–300 nm.

**Calcium Imaging**

Live-cell calcium imaging experiments followed STED imaging of FingR-PSD95-mTurq2 labeled endogenous PSD-95 puncta. Therefore, the same microscope was used. Cells were initially placed in artificial cerebrospinal fluid (ACSF, 150 mM NaCl, 5 mM KCl, 2 mM $CaCl_2$, 1 mM $MgCl_2$, 20 mM glucose and 10 mM HEPES, pH 7.35) during STED imaging. Cells were maintained at 32-35°C using a heating element and a 35 mm dish stage adaptor. In order to record mini-synaptic calcium transients (mSCTs), cells were then perfused with 10 mL of an activity- and calcium-blocking cocktail in magnesium-free ACSF (2 μM TTX, 40 μM

Nifedipine, 20 μM NBQX, 10 μM LY341495, 30 μM dantrolene, 500 μM (S)-MCPG). The pixel format was changed to 64 x 64 pixels and bidirectional scanning was activated to allow a scanning speed of ~83 Hz (12 ms frame intervals). Fields of spines were held in focus and imaged for 5-10 minutes using Adaptive Focus Control (AFC, Leica Microsystems). Following imaging of mSCTs, the cells were treated with the same inhibitor cocktail with the addition of APV (2 μM TTX, 40 μM Nifedipine, 20 μM NBQX, 10 μM LY341495, 30 μM Dantrolene, 500 μM (S)-MCPG, 100 μM APV) to verify blockade of all mSCTs.

For linescanning, the acquisition mode was changed to one-dimensional xt scanning and bidirectional scanning was turned off to achieve a scanning speed of 4000 Hz (250 μs intervals). A horizontal line was set through a spine head and its corresponding dendritic shaft such that fluorescence activity could be simultaneously recorded from both (**Fig. 2a**). Individual spines were held in focus and imaged for 10-20 minutes using AFC.

**Glutamate Imaging**

Live-cell glutamate imaging experiments followed STED imaging of FingR-PSD95-mTurq2 labeled endogenous PSD-95 puncta using the same microscope. Cells were initially placed in ACSF during STED imaging. Cells were maintained at 32-35°C using a heating element and a 35 mm dish stage adaptor. Following identification of spine PSD-95 nanomodules, the cells were perfused with 10 mL magnesium-free ACSF with 2 μM TTX in order to visualize miniature glutamate release events. The acquisition mode was then changed to one-dimensional xt-scanning in order to achieve a scanning speed of 4000 Hz (250 μs intervals). A horizontal line was set through a spine in order to record its activity. Individual spines were held in focus and imaged for 10-20 minutes using AFC.

**Data Analysis**

All analysis was performed using Fiji/ImageJ (NIH, Bethesda, MD) and OriginLabs (OriginLab Corporation, Northampton, MA). For calcium imaging analysis, custom built macros were used in ImageJ. For fieldscanning experiments of multiple spines, the macro was designed such that calcium fluorescence videos were simultaneously displayed with activity traces. Activity was displayed in both the spine head and dendrite in order to exclude events that may have originated in the dendrite due to back-propagating calcium fluctuations. A Gaussian blur (radius = 0.75) was applied to the imaging video. A region of interest (ROI) was drawn around a spine head and its corresponding dendritic shaft. A trace across the whole recording was generated and a rolling average with a window size of 6 frames or 72 ms was applied to smooth the trace data. Fluorescence was normalized using ΔF/F: $Normalized\ F = \frac{F-F_0}{F_0}$ where $F$ is the instantaneous fluorescence and $F_0$ is the fluorescence of the baseline noise. ΔF/F was applied across the entire video within both the spine and dendrite ROIs. A window of at least 500 data points of baseline noise fluorescence was selected and averaged to attain $F_0$. A threshold of at least four standard deviations above the mean noise was used to identify events using a peak finder. The events were accepted if the rise of the event started at baseline and there was no activity in the dendrite within 50 ms prior to the spine event. For events that were within a doublet or burst of activity, where the baseline could not be accurately measured, they were rejected for amplitude analysis, but they were counted for frequency analysis.

For one-dimensional xt spine recordings of calcium and glutamate activity, a modified version of the macro was utilized to accommodate for changes in acquisition mode and speed. The lines were acquired as stacks that contained two seconds of fluorescence data. The stacks were rotated 90° counterclockwise and were stitched together such that all individual lines were combined into a single image. Two ROIs were drawn on the stitched image, one around the spine head, and the other around the corresponding dendrite. The same macro was then used to generate the fluorescence trace and a rolling average with a window size of 250 data points (~60 ms) was used to smooth the data. A peak finder using a threshold of at least four standard deviations above the mean noise was used to identify peaks and display three second windows of trace activity around each event. The same criteria were used as in fieldscanning to accept and reject events. Events were normalized based on instantaneous ΔF/F. The inflection point at which the event began and a window of 100 data points (25 ms) prior to the selected inflection point was averaged to determine the $F_0$.

For kinetics analysis of individual events, events collected in ImageJ were imported into Neuromatic, a toolset developed for IgorPro (WaveMetrics, Lake Oswego, OR). An event window based on the average baseline-to-baseline temporal dynamics of the events was used to perform multiple analyses. The rise time was measured from the baseline to 80% of the peak amplitude. The decay time was measured from the event peak to 50% of the peak amplitude. The full width at half maximum was measured from the peak amplitude.

Due to a variability in baseline noise and apparent quantal size between spines, calcium and glutamate amplitudes were normalized based on minimal assumptions. Spine amplitudes were normalized based on the principle of quantal neurotransmitter release. In other words, postsynaptic responses represent unitary vesicles. Therefore, the first peak in the amplitude distribution represented the quantal size. In order to control for the variability in the first peak in spine amplitude distributions, an average bin distance from the noise amplitude distribution to the first apparent prominence in signal amplitude distribution was calculated (3 bins, 0.21 normalized GCaMP fluorescence for mSCTs; 2 bins, 0.2 normalized iGluSnFR fluorescence for mGluTs) using high frequency spines. This value was then applied to every spine based on the noise for each spine. This way all spines were normalized the same way based on the all of acquired data.

In order to perform Poisson analysis all event amplitude data were plotted as histograms in OriginLabs. Data were then fit using the Poisson Function (P): $P(X = x) = N\frac{\lambda^x e^{-\lambda}}{x!}$ where $N$ is a scaling factor, because the Poisson function applies to fractional probabilities <1, $x$ is bin number, and $\lambda$ is mean quantal amplitude. Multiple Poisson functions were modeled based on the principle of quantal neurotransmitter release. The quantal size of each distribution was determined to be the unitary postsynaptic response and was based on the position of the first prominence in the distribution. The subsequent quanta were modeled based on even-spacing from one another, such that they represent linear summations of the unitary event. Single-term, two-term, three-term, and four-term Poisson functions were fit over all data sets. Multiple Poisson Functions were generated by adding the number of Poisson terms into a single formula. The Multiple Poisson Function fit the data to a single cumulative curve. The best fit models compared by Akaike Information Criteron were used to represent the data.

**Image Processing**

The calcium imaging data was minimally processed. A Gaussian blur (radius = 0.75) was applied to field recorded calcium images, and the fluorescence intensity of the ROIs selected for analysis were normalized by $\Delta F/F$. Image processing for FingR-PSD95-mTurq2 involved first applying a Fast Fourier Transform (FFT) filter to remove detector background noise. Following FFT filtering a Gaussian blur (radius = 0.8-1.5) was applied to the images.

**Immunocytochemistry**

Immunostaining was performed for FingR-PSD95-mTurq2 validation experiments. Cells were taken out of the incubator at DIV 25-27. The media was removed and replaced with 4% PFA and 2% sucrose with 0.000375% glutaraldehyde (cat #: 16000, Electron Microscopy Science Hatfield, PA) for eight minutes. After eight minutes, the cells were washed one time with 1x Phosphate Buffered Saline (PBS) solution. The PBS was replaced with 1 mg/mL sodium borohydride (cat #: 213462-25 g, Millipore Sigma) and the cells were placed at 4°C for 15 minutes. After 15 minutes, the cells were washed three times with 1x PBS and were placed in 0.2% cold water fish gelatin (cat #: G7041-100G, Millipore sigma) and 1% ovalbumin (cat #: A5503, Millipore Sigma) containing blocking buffer with 0.01% saponin (cat #: 47036, Millipore sigma), to permeablize cells, for at least two hours at room temperature or over-night at 4°C. After incubation, the blocking buffer was replaced by blocking buffer containing primary antibodies and 0.01% saponin. Antibodies against Bassoon, PSD-95 and GFP (which can be used to detect FingR-PSD95-mTurq2) were used. The cells were incubated in primary antibodies for two hours at room temperature or over-night at 4°C. After incubation, the cells were washed three times with PBS.

**Pharmacologic Agents**

The following drugs were used to collect miniature synaptic calcium transients and miniature glutamate transients: tetrodotoxin-citrate (TTX, cat #: 1069, Tocris Bioscience, Bristol, United Kingdom), nifedipine (cat #: 1075, Tocris Bioscience, Bristol, United Kingdom), 2,3-dioxo-6-nitro-7-sulfamoyl-benzo[f]quinoxaline (NBQX, cat #: 0373, Tocris Bioscience, Bristol, United Kingdom), LY341495 (cat #: 1209, Tocris Bioscience, Bristol, United Kingdom), dantrolene (cat #: 0507, Tocris Bioscience, Bristol, United Kingdom), (S)-α-methyl-4-carboxyphenylglycine ((S)-MCPG, cat #: HB0056 Hello Bio Inc., Princeton, NJ), D-(2R)-amino-5-phosphonopentanoate (D-AP5, cat #: 0106, Tocris Bioscience, Bristol, United Kingdom).

**Antibodies**

Primary antibodies: mouse monoclonal IgG1 anti-PSD-95 clone 7E3-1B8 (1:200, cat #: MA1-046, Thermo Fisher Scientific), rabbit polyclonal anti-GFP (1:2000, cat #: ab290 Abcam, Cambridge, MA), guinea pig monoclonal recombinant IgG anti-Bassoon (1:300, cat #: 141 318, Synaptic Systems, Gottingen, Germany). Secondary antibodies: goat anti-mouse IgG1 Atto-647N (1:500, cat#: 610-156-040, Rockland Inc, Limerick, PA), donkey anti-guinea pig AlexaFluor-594 (1:500, cat #: 706-586-148, Jackson ImmunoResearch, West Grove, PA), donkey anti-rabbit AlexaFluor-488 (1:500, cat #: 711-545-152, Jackson ImmunoResearch)

**Statistics and reproducibility**

Statistical analyses were done using GraphPad Prism (GraphPad Software, Boston, Massachusetts) and OriginLab. Data are expressed as means with individual values overlaid as dots. In violin plots the median and 25$^{th}$ and 75$^{th}$ quartiles are shown. Error bars represent standard error of the mean (SEM). Statistical significance for APV treatment and photobleaching was performed using paired t-test. Differences among cumulative probability distributions was determined by K-S nonparametric tests. Full width at half maximum, rise time, and decay time between groups were compared using one-way ANOVA with Tukey's posthoc test. Raw peak event amplitudes from one quanta events and mean event frequency between groups were compared using unpaired t-test. Statistically significant results were determined with p values < 0.05, with * representing $p < 0.05$, ** representing $p < 0.005$, *** representing $p < 0.0005$, and **** representing $p < 0.0001$. Data were collected from a minimum of three neurons from three independent transfections and neuronal preps.
